# Lemur: A Single-Cell Foundation Model with Fine-Tuning-Free Hierarchical Cell-Type Generation for *Drosophila melanogaster*

**DOI:** 10.1101/2025.02.04.636468

**Authors:** Jorge Botas, Johnathan Jia, Sasidhar Pasupuleti, Hu Chen, Xia Hu, Zhaozhuo Xu, Zhandong Liu

## Abstract

Single-cell genomics has revolutionized our understanding of cellular heterogeneity, but automating its analysis remains an open challenge. Cell-type annotation represents a critical bottleneck, particularly as datasets grow in size and complexity. While foundation models have shown promise in addressing this challenge, existing approaches require extensive fine-tuning for effective cell-type annotation. Here, we present Lemur (<u>L</u>arge <u>E</u>xpression <u>M</u>odel for <u>U</u>nderstanding sc<u>R</u>NA-seq), a single-cell foundation model specifically designed for *Drosophila melanogaster*. Lemur achieves fine-tuning-free cell-type annotation through comprehensive pre-training on an integrated whole-organism atlas with a unified cell-type annotation schema. To leverage this unified schema, we developed a dedicated hierarchical cell-type decoder architecture. This approach enables Lemur to generate consistent cell-type predictions across multiple levels of granularity without requiring additional training on new datasets. The model demonstrates strong performance across diverse tissue types, experimental conditions, and sequencing technologies. It also achieves batch-effect correction without explicitly training for this task. This automated analysis capability positions Lemur as an effective tool for the fly research community. Beyond its immediate applications, Lemur establishes a framework for accelerating biological discovery. It enables rapid iteration between computational predictions and experimental validation in the highly controlled *Drosophila melanogaster* system, with potential implications for translational research in human biology, particularly in aging and neurodegenerative disease studies.

---

The exponential growth of single-cell genomics data has revolutionized our understanding of cellular heterogeneity while creating new computational challenges for data analysis. Traditional analysis pipelines involve a series of complex steps—including quality control, batch correction, dimensionality reduction, clustering, and cell-type annotation—each requiring specialized tools and expertise. Among these steps, cell-type annotation has remained a critical bottleneck, particularly as datasets continue to expand in size and complexity. The traditional approach relies on manual curation by domain experts who must meticulously evaluate gene expression patterns to assign cell-type labels, a process that is not only time-consuming but also limits reproducibility^[1]^. This expert-dependent methodology faces several significant challenges: inconsistent interpretation of marker genes across experts, varying nomenclature conventions, and divergent classification criteria across laboratories and studies.

Foundation models, originally popularized in natural language processing and computer vision, have emerged as a promising solution to these challenges. These large-scale deep learning models, pre-trained on massive, heterogeneous datasets, learn versatile representations that can be adapted to a broad range of downstream tasks^[2]^. This concept has been successfully extended to single-cell genomics through single-cell foundation models (scFMs)^[3]^, which leverage transformer-based architectures to capture complex gene-gene relationships and cell-level variations in a self-supervised manner. Recent advances in human scFMs, such as Geneformer ^[4]^ and scGPT ^[5]^, have demonstrated remarkable success in various tasks including cell-type annotation, batch effect correction, and perturbation response prediction.

Developing a species-specific foundation model using *Drosophila melanogaster* offers a unique opportunity to overcome these limitations through its highly controlled experimental environment. Unlike human studies, fly research benefits from standardized genetic backgrounds and laboratory conditions, minimizing confounding variables that complicate data interpretation. Most notably, the recent development of comprehensive single-nucleus atlases spanning the entire organism—including the Fly Cell Atlas^[6]^, Aging Fly Cell Atlas^[7]^, and Alzheimer’s Disease Fly Cell Atlas^[8]^—provides an unprecedented resource with unified cell-type annotations and minimal technical variation. While adapting human models to fly data is impractical due to complex evolutionary relationships and often ambiguous gene orthology^[9]^, this integrated data framework enables the development of dedicated fly-specific foundation models capable of extracting genuine biological patterns with greater fidelity.

Beyond its advantages for model development, *Drosophila melanogaster* represents an ideal system for bridging computational predictions and experimental validation. The fly’s genetic tractability, facilitated by sophisticated tools for precise genetic manipulation, combined with its short life cycle enables high-throughput validation of model-derived insights. This creates an efficient feedback loop between *in silico* discoveries and *in vivo* verification, accelerating the pace of biological discovery. Importantly, despite the evolutionary distance necessitating species-specific computational approaches, the conservation of core biological pathways between flies and humans—particularly in aging and disease processes—positions fly-based models as valuable tools for translational research. This is evidenced by numerous break-through discoveries in neuronal development, aging, and neurodegenerative diseases that have been successfully translated to human biology^[10]^.

Here, we present Lemur (Large Expression Model for Understanding scRNA-seq), a single-cell foundation model tailored to *Drosophila melanogaster* that achieves true fine-tuning-free cell-type annotation through the synergy of comprehensive pretraining data and architectural innovation. At its core, Lemur leverages the most comprehensive whole-organism snRNA-seq dataset available for the adult fly^[8]^, encompassing over 624,458 cells spanning 219 cell types. Crucially, this dataset provides a unified cell-type annotation schema derived from the integration of multiple atlases^[6, 7, 11]^, enabling the development of a novel transformer architecture that capitalizes on its hierarchical organization. Lemur’s innovative design combines a transcriptomic encoder with a dedicated hierarchical cell-type decoder, which generates cell-type labels in a consistent progression from broad categories to specific subtypes. This approach ensures biological coherence across different levels of annotation granularity, facilitating reliable cell-type identification even in complex tissues where fine-grained annotations may be challenging to determine with high confidence. Most notably, Lemur can accurately assign cell-type labels to unseen data without requiring additional training, a breakthrough we term “fine-tuning-free annotation”. This capacity stands in contrast to existing human scFMs, which often demand a specialized fine-tuning phase to handle new datasets. Furthermore, Lemur demonstrates robust performance across multiple tissue types, successfully bridges different sequencing modalities, and effectively addresses batch-effect correction in a zero-shot setting. Together, these capabilities position Lemur as a powerful tool for the fly research community, enabling automated and scalable analysis of single-cell data to accelerate discoveries in developmental biology, aging, and neurodegenerative disease research.

## Results

### Model Overview

Lemur introduces a novel architecture specifically designed to leverage the unique advantages of *Drosophila* single-cell data (**Figure 1**). The model was pre-trained on the most comprehensive and up-to-date single-nucleus RNA-seq dataset available for the adult fly^[8]^, comprising over 624,458 cells expressing 16,195 genes. This dataset represents a significant advancement over existing resources in regards to the number of cell types by integrating annotations from multiple atlases into a unified schema, including the Fly Cell Atlas^[6]^, Aging Fly Cell Atlas^[7]^, and a comprehensive optic lobe atlas^[11]^. The t-distributed stochastic neighbor embedding (t-SNE) visualization of the pretraining corpus depicts distinct clusters corresponding to 19 major cell types including neurons, glia, muscle cells, and other tissue-specific populations (**Figure 1A**).

**Figure 1:**
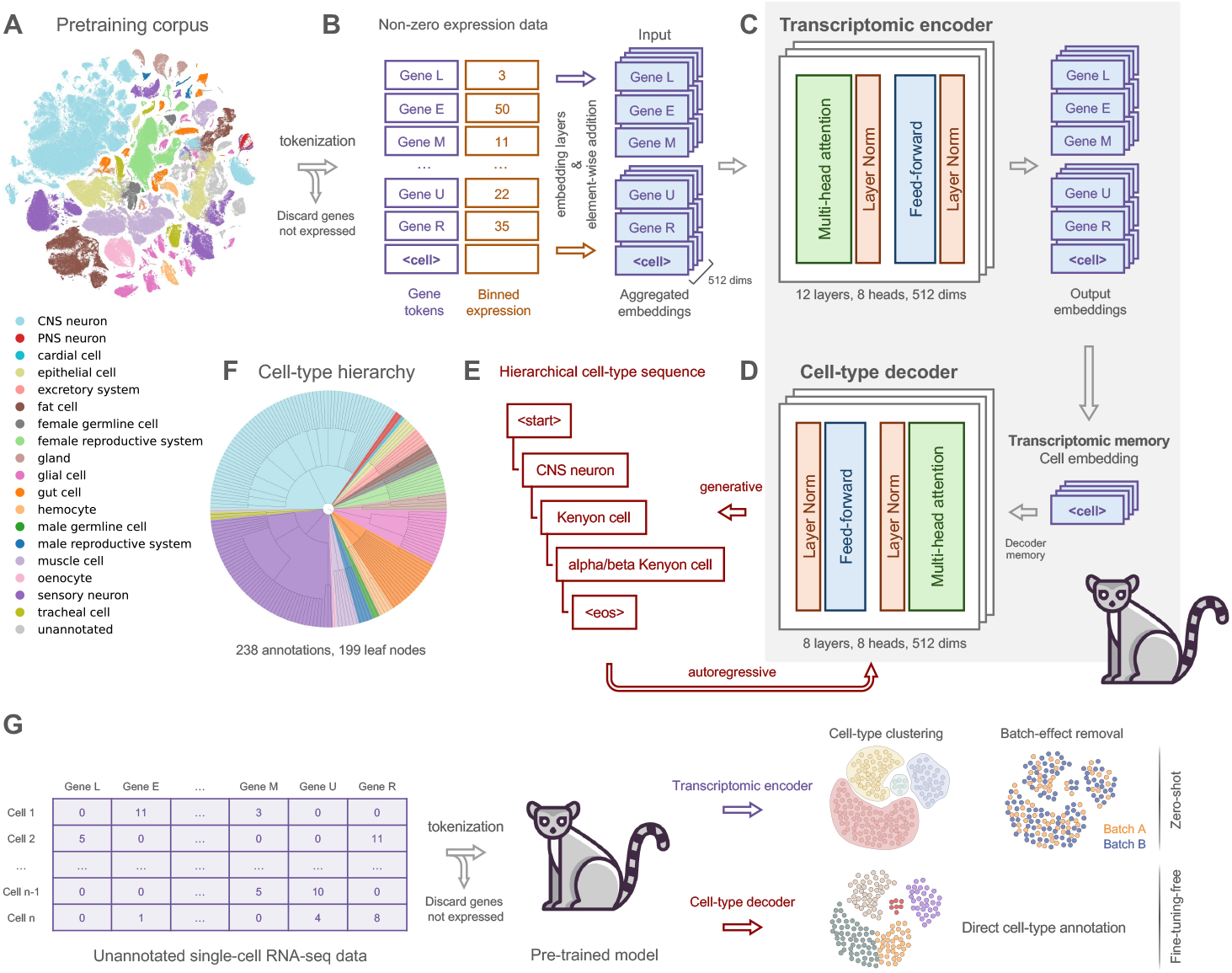
Lemur’s architecture and workflow. **(A)** t-SNE visualization of the pretraining corpus prior to model training, displaying distinct clusters of 19 major cell types including neurons, glia, muscle cells, and other tissue-specific cell populations. **(B)** Data preprocessing pipeline that converts single-cell transcriptomes into model inputs. For each cell, only genes with non-zero expression values are processed. These expression values are discretized through rank-based binning across each cell’s transcriptome. Each gene and its binned expression value are tokenized separately and embed-ded, followed by aggregation and prepending of a `cell` token that captures the cell’s overall expression profile. **(C)** Architecture of the transcriptomic encoder comprising 12 transformer layers (512 dimensions, 8 attention heads). Each layer contains multi-head self-attention and feed-forward modules with layer normalization, processing the aggregated embeddings to generate context-aware gene and cell representations. **(D)** Lemur’s novel hierarchical cell-type decoder (8 layers, 512 dimensions, 8 attention heads) uses the `cell` embedding output by the encoder as a transcriptomic memory. **(E)** The decoder generates cell-type labels in an autoregressive manner, starting with `start` and progressively refining predictions from broad categories (e.g., CNS neuron) to specific cell types (e.g., alpha/beta Kenyon cell), ending with an end-of-sentence token `eos`. **(F)** Comprehensive cell-type hierarchy comprising 238 distinct annotations organized in a tree structure with 199 leaf nodes. This hierarchical organization enables biologically meaningful predictions at multiple levels of granularity. **(G)** Overview of Lemur’s inference workflow. The pre-trained model processes new single-cell data and generates two types of outputs: 1) transcriptomic encoder embeddings that enable zero-shot cell-type clustering and batch effect correction, useful for further downstream analyses and visualization; and 2) hierarchical cell-type annotations from the dedicated decoder that provides multiple levels of annotation resolution without any fine-tuning requirement (fine-tuning-free).

The model’s architecture consists of two primary components: a transcriptomic encoder and a hierarchical cell-type decoder. To process input data, Lemur employs a specialized preprocessing pipeline that focuses on expressed genes—those with non-zero expression values (**Figure 1B**). For each gene, the expression level undergoes rank-based binning to create discrete categories, providing a robust representation that minimizes technical variations^[5]^. These gene-expression pairs are then independently embedded and aggregated, with a prepended `cell` token that serves to capture the cell’s overall expression profile.

The transcriptomic encoder comprises twelve transformer layers, each containing multi-head self-attention and feed-forward modules with layer normalization (**Figure 1C**). Operating with 512 dimensions and eight attention heads, these layers process the aggregated embeddings to generate context-aware gene and cell representations. This deep architecture enables the model to capture complex relationships between genes and their expression patterns within each cell.

Lemur introduces an architectural innovation through its hierarchical cell-type decoder (**Figure 1D**). This decoder, consisting of eight transformer layers with matching dimensionality to the encoder, utilizes the `cell` embedding as a transcriptomic memory to generate cell-type labels in an autoregressive manner. Starting with a `start` token, the decoder progressively refines its predictions from broad categories to specific cell types (e.g., from CNS neuron to alpha/beta Kenyon cell), concluding with an end-of-sentence `eos` token (**Figure 1E**). This approach ensures consistency across different levels of annotation granularity, particularly valuable in complex tissues where fine-grained annotations may be challenging to determine with high confidence.

The design of Lemur’s architecture was motivated by the harmonized nature of our pretraining data, which enabled the natural extraction of a comprehensive cell-type ontology. This ontology comprises a total of 238 annotations organized in a tree structure with 199 leaf nodes (**Figure 1F, Supplementary Figure 1**), providing a robust framework for the model’s hierarchical predictions. This structured approach to cell-type annotation represents a significant advance over existing methods, enabling more nuanced and biologically meaningful analysis of cellular heterogeneity.

After pretraining, Lemur can process new single-cell RNA-seq data through two complementary pathways (**Figure 1G**). First, the transcriptomic encoder generates cell embeddings that enable zero-shot applications such as cell-type clustering and batch effect correction. Second, the hierarchical decoder provides multi-resolution cell-type annotations without requiring any fine-tuning, a capability we term “fine-tuning-free” annotation. This dual output structure makes Lemur a versatile tool for analyzing new datasets, combining the advantages of unsupervised representation learning with precise cell-type classification.

Ontology reconstructed and visualized using ETE Toolkit^[12]^.

### Lemur Enables Fine-Tuning-Free Cell-Type Annotation Through Hierarchical Decoding

#### Generative Cell-Type Annotation

Lemur’s decoder implements a novel generative approach to cell-type annotation by progressively refining its predictions through a hierarchical structure. We evaluated the model’s performance on an independent single-cell RNA-seq dataset from the adult fly brain ^[13]^, chosen for its cellular and transcriptomic complexity. **Figure 2A** illustrates the step-by-step annotation process for CNS neurons in the adult fly brain. Starting from unannotated data, the decoder initially assigns a broad category (i.e., CNS neuron) and then incrementally refines its prediction through intermediate categories (e.g., medulla intrinsic, distal medullary amacrine) until arriving at specific cell types (e.g., medulla intrinsic neuron Mi4, distal medullary amacrine neuron Dm3). This generative progression mirrors the natural organization of cell types in our unified annotation schema, thereby ensuring biological coherence across all prediction levels.

**Figure 2:**
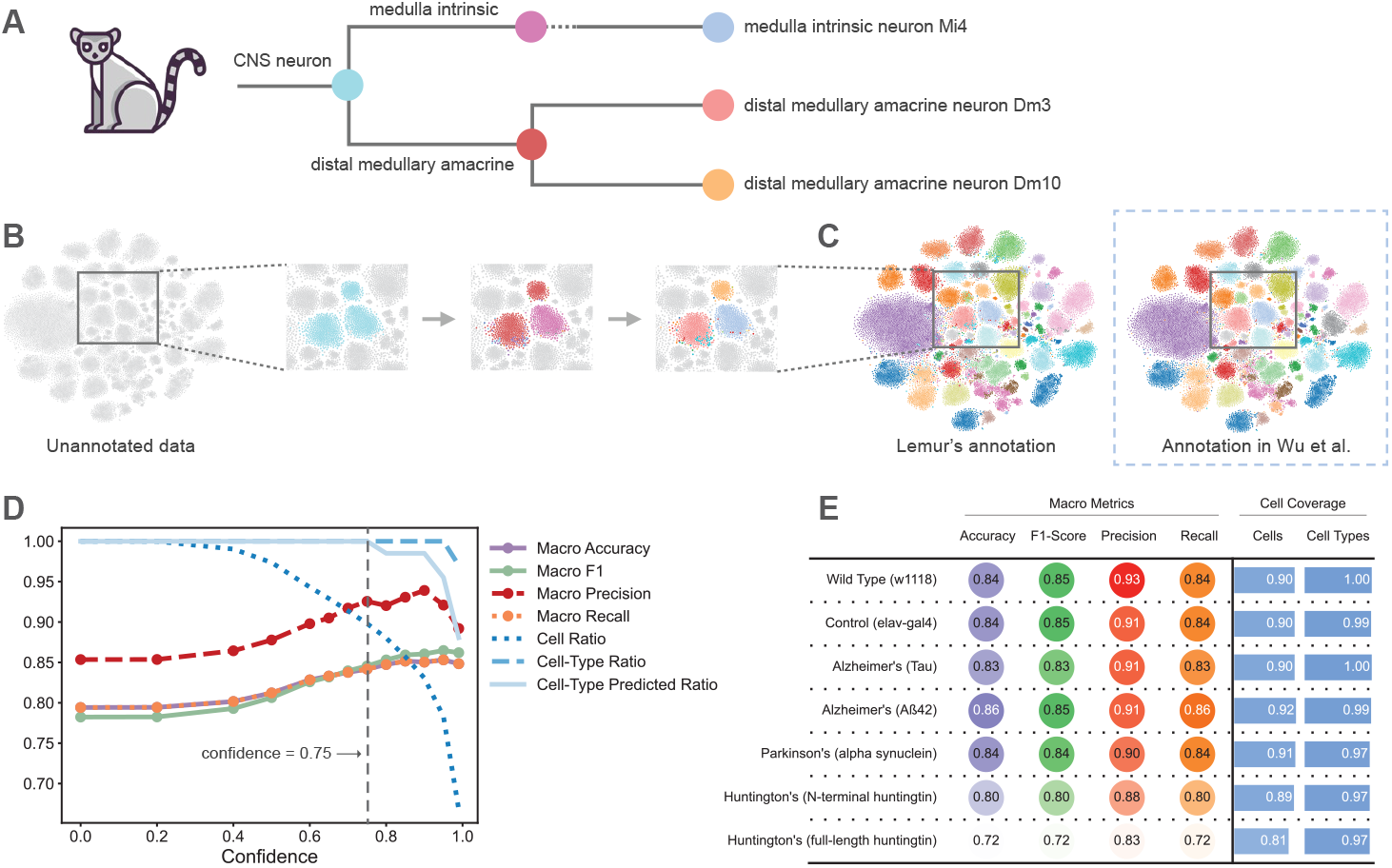
Lemur Hierarchical Cell-Type Annotation and Performance Across Conditions. **(A)** Illustration of Lemur’s hierarchical decoding process, showing the progressive refinement from broad categories (e.g., CNS neuron) to specific cell types (e.g., medulla intrinsic neuron Mi4, distal medullary amacrine neurons Dm3 and Dm10). **(B)** t-SNE visualization of the annotation process on unannotated brain data, with sequential panels demonstrating the stepwise resolution of cellular identities. **(C)** Comparison of high-resolution annotations (67 distinct cell types) between Lemur’s predictions (left) and expert annotations (right) in t-SNE space, revealing strong concordance. **(D)** Performance metrics as a function of confidence threshold, including macro accuracy, F1-score, precision, recall, and retention ratios for cells and cell types; the vertical dashed line indicates the optimal threshold of 0.75. **(E)** Model performance across wild-type and various disease model datasets, showcasing macro metrics (accuracy, F1-score, precision, recall) alongside cell coverage statistics. Robust performance is maintained across conditions, with high precision and comprehensive cell-type coverage.

The hierarchical annotation process is further visualized through t-SNE projections in **Figure 2B**, which demonstrate how cellular identities are progressively resolved from the unannotated input. At the highest resolution (**Figure 2C**), Lemur accurately identified all 67 distinct cell types present in the dataset, exhibiting remarkable concordance with expert annotations.

#### Confidence-Based Performance Optimization

To optimize performance, we systematically examined the relationship between the model’s confidence and its predictive accuracy (**Figure 2D**). Our analysis revealed that even without applying a confidence filter, the model achieves a strong baseline performance with macro accuracy and F1-scores of 79%. As the confidence threshold increases, improvements in precision and accuracy become evident, with an optimal threshold identified at 0.75. At this level, the model retains 90% coverage of cells while preserving all annotated cell types, as demonstrated by the cell ratio and cell-type ratio curves.

This balance between prediction reliability and comprehensive cellular coverage underscores the robustness of Lemur’s annotation strategy.

#### Robust Performance Across Disease States

To further assess Lemur’s applicability, we evaluated its performance across multiple neurodegenerative disease models (**Figure 2E**). The model demonstrated robust performance across all conditions, achieving accuracies of 83–86% for Alzheimer’s disease samples (Tau and *Aβ*42 models), 84% for Parkinson’s disease (alpha-synuclein), and 72–80% for Huntington’s disease models (N-terminal and full-length huntingtin). Notably, high precision (0.88–0.93) and comprehensive cell-type coverage (0.97–1.00) were maintained across most conditions. This consistent performance is particularly striking given that Lemur was trained on single-nucleus RNA-seq data yet performs effectively on single-cell RNA-seq data. Such robust cross-modality and cross-condition transfer suggest that Lemur has captured fundamental cell-type-specific expression patterns that generalize beyond technical variations.

### Zero-Shot Batch-Effect Correction While Maintaining Biological Integrity

A critical challenge in single-cell analysis is the removal of systematic technical variations between experimental batches while preserving genuine biological differences. To evaluate Lemur’s capacity for zero-shot batch effect correction without fine-tuning, we analyzed single-cell RNA sequencing data from leg and wing tissues^[6]^ containing four distinct experimental batches. Initial visualization of the unintegrated data revealed cells clustering primarily by batch (**Figure 3A**, top), with technical variation dominating the underlying biological structure of six distinct cell populations (**Figure 3A**, bottom). After processing through Lemur’s transcriptomic encoder, cells from different batches showed effective mixing (**Figure 3B**, top) while enhancing the separation of distinct cell populations (**Figure 3B**, bottom), despite the model never being explicitly trained for batch correction.

**Figure 3:**
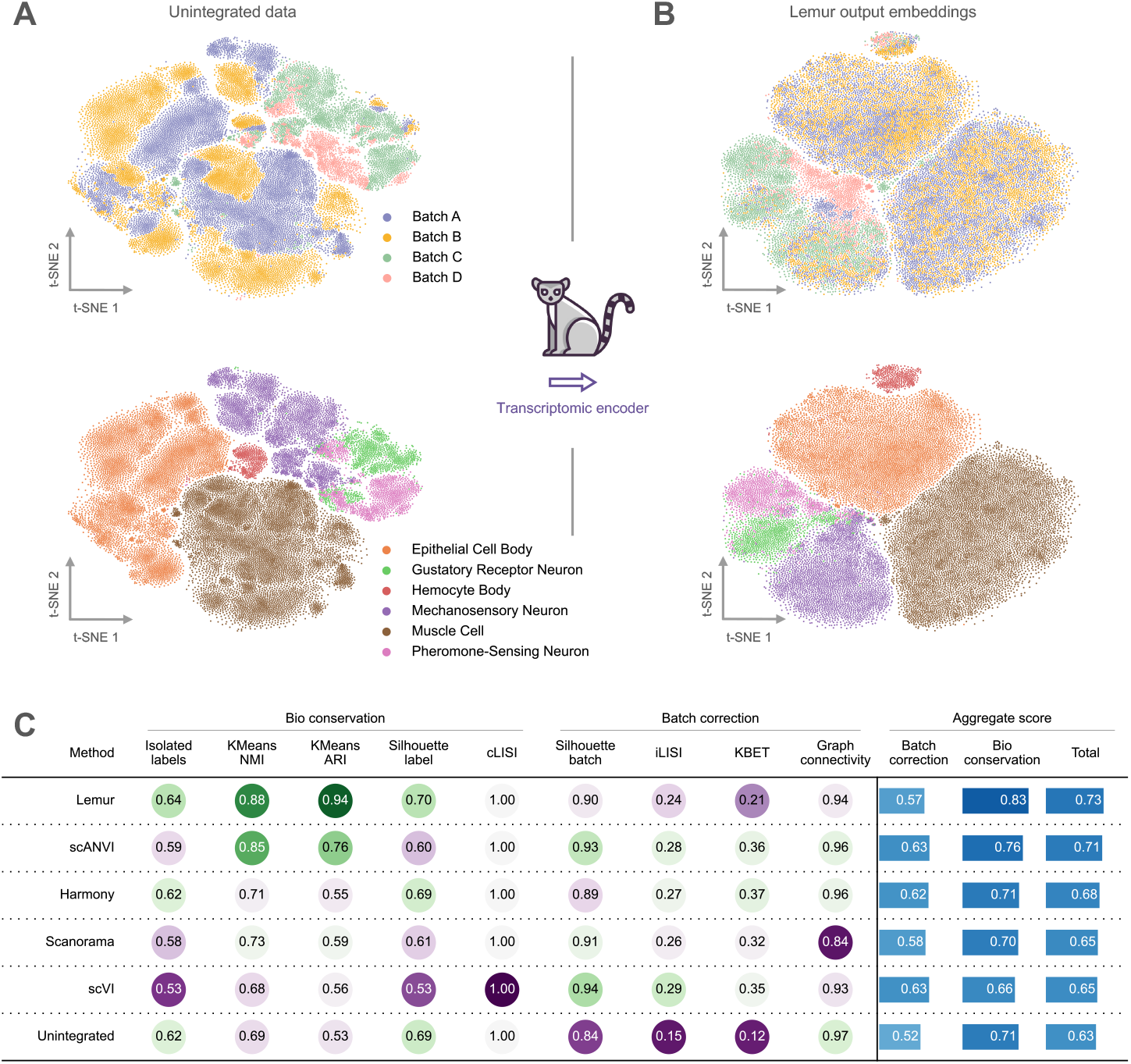
Lemur effectively removes batch effects while preserving biological signals in a zero-shot manner. **(A)** t-SNE visualization of unintegrated single-cell RNA sequencing data colored by experimental batch (top) and cell type (bottom). Cells cluster predominantly by technical batch, partially obscuring the biological relationships between the six distinct cell populations. **(B)** t-SNE visualization of Lemur’s output embeddings demonstrating effective batch integration. Cells from different experimental batches are well-mixed (top) while biological distinctions between cell types become more prominent (bottom). Lemur achieves this integration without training for batch effect correction or finetuning on this dataset. **(C)** Quantitative comparison of integration performance across methods. Metrics are grouped into biological conservation (measuring preservation of cell-type structure) and batch correction (measuring removal of technical variation). Each metric is scaled from 0 to 1, where higher values indicate better performance. Circle color represents relative performance (normalized by column, for enhanced contrast). Aggregate scores are computed as weighted averages of individual metrics. Despite not having an explicit batch correction objective, Lemur (aggregate score: 0.73) outperforms specialized batch correction methods, including scANVI (0.71), which uses supervised celltype information. Notably, Lemur achieves superior biological signal preservation (0.83) while maintaining effective batch correction (0.57), demonstrating that foundation model pre-training can implicitly learn to recognize and correct technical variation.

To quantitatively assess integration performance, we employed established metrics that evaluate both the correction of batch effects and the conservation of biological signal ^[14]^. For batch correction, we considered the silhouette batch score ^[15]^, integration Local Inverse Simpson’s Index (iLISI) ^[16]^, k-Batch-Effect Test (kBET) ^[17]^, and graph connectivity ^[18]^. For biological conservation, we measured cluster preservation using KMeans Normalized Mutual Information (NMI) ^[19]^, Adjusted Rand Index (ARI) ^[20]^, cell-type silhouette scores, and conservation Local Inverse Simpson’s Index (cLISI) ^[16]^.

A comprehensive comparison against state-of-the-art integration methods underscores Lemur’s superior performance (**Figure 3C**). Lemur achieves an overall batch correction score of 0.57 and a biological conservation score of 0.83, yielding an aggregate performance score of 0.73—the highest among all methods tested. Notably, Lemur performance matches and even surpasses specialized integration algorithms such as scANVI^[21]^ (aggregate score 0.71), which require additional fine-tuning with cell-type annotations and batch information. The model further excels in preserving rare cell populations (isolated labels score: 0.64) and attains the highest KMeans ARI (0.94) and NMI (0.88) scores, surpassing dedicated methods like Harmony^[16]^ (ARI: 0.55, NMI: 0.71), scVI^[22]^ (ARI: 0.56, NMI: 0.68) and Scanorama^[23]^ (ARI: 0.58, NMI: 0.73).

Together, these findings indicate that Lemur’s pre-training on the *Drosophila melanogaster* transcriptome is a onetime effort that endows it with the intrinsic ability to recognize and correct technical variations while faithfully preserving biological signals. This emergent capability, achieved without explicit batch correction objectives, highlights the potential of large-scale self-supervised learning to capture fundamental principles of transcriptional variation that generalize across diverse experimental conditions.

## Discussion

Lemur represents a major advancement in single-cell genomics analysis by providing a fine-tuning-free approach to both cell-type annotation and batch effect correction. By eliminating the need for extensive model adaptation, Lemur allows researchers to move directly from raw data to biological insights, thereby streamlining the analytical pipeline and reducing technical barriers.

The immediate usability of Lemur is rooted in its innovative architecture and its comprehensive pretraining on an integrated *Drosophila melanogaster* atlas with a unified cell-type annotation ontology. Unlike many existing single-cell foundation models that require dataset-specific fine-tuning, Lemur can be deployed directly on new datasets while maintaining high accuracy across diverse tissue types and experimental conditions. This “ready-to-use” nature democratizes access to advanced computational methods, making sophisticated cell-type annotation accessible even to researchers with limited computational expertise.

A key feature of Lemur is its hierarchical decoder, which enhances annotation accuracy by providing biologically meaningful predictions at multiple levels of granularity. Researchers can explore their data from broad cell categories down to specific subtypes, confident that the model’s predictions remain consistent and informative across these resolutions. Moreover, the model’s robust performance across challenging tissue types and disease states underscores its strong generalization capabilities.

The synergy between Lemur and the *Drosophila* model system creates a powerful platform for biological discovery. The standardized genetic backgrounds and controlled laboratory conditions in fly research, combined with Lemur’s accurate *in silico* predictions, pave the way for an efficient cycle of computational prediction and experimental validation. This integration is especially valuable for studies in developmental biology, aging, and neurodegenerative diseases, where rapid iterations between prediction and verification can greatly accelerate research progress.

Despite these strengths, two potential limitations warrant consideration. First, because Lemur is pretrained exclusively on *Drosophila melanogaster* data, its direct applicability to other organisms may be limited. While the underlying design principles are broadly relevant, adapting the model for use in other species might require additional training or validation efforts. Second, although Lemur is able to map new cell types within its preexisting hierarchical framework, it does not perform true zero-shot annotation. That is, while the model provides hints of broader resolution by integrating novel cell types into its established hierarchy, fully characterizing cell-type classes that were never seen during pre-training remains an open challenge in the field.

In conclusion, Lemur offers a streamlined solution that alleviates many of the technical hurdles traditionally associated with single-cell analysis. By enabling accurate cell-type annotation and robust batch effect correction without the need for extensive fine-tuning, Lemur empowers researchers to concentrate on unraveling complex biological questions rather than troubleshooting computational challenges. With its open-source release and broad accessibility, this tool is well positioned to benefit the broader research community. This accessibility, combined with the model’s performance across diverse experimental conditions, positions Lemur as a catalyst for accelerating discoveries in aging research and neurodegenerative disease studies. By bridging the gap between computational prediction and experimental validation in *Drosophila melanogaster*, Lemur not only streamlines current research workflows but also opens new possibilities for understanding cellular complexity in health and disease.

## Methods and Materials

### Dataset and Preprocessing

#### Training Data

Lemur was pre-trained on the most comprehensive and up-to-date single-nucleus RNA-seq dataset available for the adult fly^[8]^, comprising 624,458 cells and spanning 16,195 genes. This whole-organism dataset encompasses both whole head and body tissues, with annotations for 219 distinct cell types (143 labels from head, 74 from body). When hierarchically organized, a total of 238 annotations are considered across resolutions, with 199 terminal nodes and 19 top-level categories. The dataset represents a significant advancement in unified cell-type annotation, integrating data from the Fly Cell Atlas^[6]^, Aging Fly Cell Atlas^[7]^, and comprehensive optic lobe atlas^[11]^.

To ensure data quality, we implemented rigorous preprocessing steps including the removal of low-quality cells with mitochondrial gene expression exceeding 2%. The unified annotation schema was carefully preserved across all cells, maintaining consistent labeling from broad categories to specific cell types. Data loading and preprocessing were facilitated by the datasets library from Hugging Face, enabling efficient handling of the large-scale dataset during training.

#### Evaluation Datasets

To rigorously evaluate Lemur’s performance and generalization capabilities, we employed multiple independent test datasets spanning different tissues and disease conditions. For central nervous system evaluation, we utilized wild-type brain and Tau disease model datasets from ^[13]^, which are publicly accessible through the Alzheimer’s Disease Knowledge Portal (Synapse:syn35798807.2). Additional brain datasets from various disease models, developed at Baylor College of Medicine, were used to assess the model’s robustness across pathological conditions. These datasets are currently under review or part of manuscripts in preparation. For peripheral tissue evaluation, we leveraged leg and wing datasets generated by the Kristin Scott laboratory at University of California, Berkeley ^[6]^. These datasets, along with additional tissue-specific data, are publicly available through the Fly Cell Atlas portal. The diverse nature of these evaluation datasets, encompassing both neural and non-neural tissues across healthy and disease states, enabled comprehensive assessment of Lemur’s fine-tuning-free annotation capabilities and batch effect correction performance.

### Input Processing and Embedding

The model processes single-cell transcriptomes through a specialized tokenization scheme that focuses on non-zero expressed genes, with a maximum input sequence length of 2,048 genes. This architectural choice accommodates 99.80% of the cellular transcriptomes in our dataset without truncation (**Supplementary Figure 2**), ensuring comprehensive coverage of gene expression patterns while maintaining computational efficiency. For each cell, we consider the set of expressed genes 𝒢_*e*_ = {*g* : *g*(*x*) > 0} and their corresponding expression values. Expression values are transformed through rank-based binning within each cell:

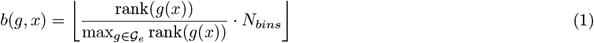

where *N*_*bins*_ represents the number of discrete expression bins. This binning strategy creates a robust, nonparametric representation that is less sensitive to technical variations and batch effects.

Each gene token and its binned expression value are embedded independently through separate embedding layers *E*_*gene*_ and *E*_*expr*_, then combined through element-wise addition:

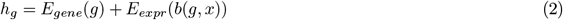

A special `cell` token is prepended to each sequence, serving as an aggregator for the cell’s overall transcriptional state through the encoder’s self-attention mechanism.

### Model Architecture

Lemur introduces a novel expression-to-annotation encoder-decoder transformer architecture designed to leverage the unique advantages of unified Drosophila single-cell data. The architecture consists of two primary components: a transcriptomic encoder and a cell-type decoder, both sharing a hidden dimension of 512 to enable efficient information flow. Throughout the model, we apply a dropout ratio of 0.1 in self-attention and fully connected layers to prevent overfitting, with layer normalization (*ϵ* = 1 *×* 10^−5^) ensuring stable training.

### Transcriptomic Encoder

The encoder comprises 12 transformer layers, each containing an eight-head self-attention module followed by a feedforward neural network. The self-attention mechanism employs scaled dot-product attention with a dimensionality *d*_*k*_ of 64 per head (512 total):

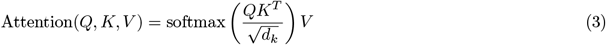

The feed-forward network expands the hidden dimension to 2048 through two linear transformations with a Gaussian Error Linear Unit (GELU) activation function between them:

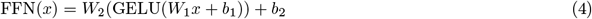

Each transformer layer applies layer normalization before both the attention and feed-forward components, following the pre-norm architecture. Notably, the encoder operates without positional encodings, as transcriptomic data lacks inherent sequential order.

### Cell-Type Hierarchy

The cell-type hierarchy ℋ = (*V, E*) emerges naturally from our unified annotation schema, where *V* represents the set of cell-type terms and *E* represents the implicit parent-child relationships. During training, we preserve these hierarchical relationships in the sequence of annotations for each cell 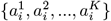, where annotations progress from broad to specific categories.

The complete hierarchy comprises 238 distinct annotations organized in a tree structure with 199 leaf nodes. This organization provides a natural framework for the decoder to learn the progression from broad cell-type categories to specific subtypes.

### Hierarchical Cell-Type Decoder

The cell-type decoder represents a significant architectural innovation in Lemur, consisting of 8 specialized transformer layers that combine hierarchical sequence generation with transcriptomic memory retention. Unlike traditional classification approaches that treat cell-type annotation as independent decisions at each resolution, our decoder generates a coherent sequence of cell-type labels that naturally progress from broad to specific categories.

#### Architecture Overview

Each decoder layer integrates three key components: masked self-attention for autoregressive generation, cross-attention to the cell’s transcriptomic state, and a feed-forward network for transformation. The layer operations can be expressed as:

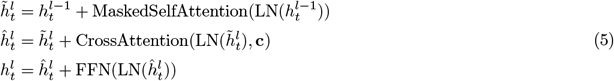

where LN denotes layer normalization and 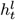 represents the hidden state at position *t* in layer *l*.

#### Transcriptomic Memory Mechanism

The decoder maintains a persistent connection to the cell’s transcriptional profile through its cross-attention mechanism. Given the cell embedding **c** from the encoder, the cross-attention computation is:

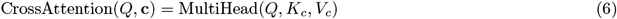

where *K*_*c*_ = *W*_*K*_**c** and *V*_*c*_ = *W*_*V*_ **c** are learned projections of the cell embedding. This mechanism allows the decoder to ground each hierarchical prediction in the cell’s underlying transcriptional state, ensuring biological relevance across all levels of annotation.

#### Hierarchical Token Generation

The decoder generates cell-type labels autoregressively, with each prediction *y*_*t*_ conditioned on both previous tokens and the transcriptomic memory:

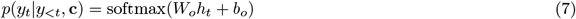

Crucial to the hierarchical generation is the incorporation of sinusoidal position embeddings that encode the sequential nature of the annotation hierarchy:

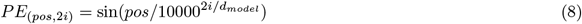

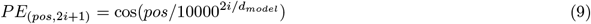

These embeddings are added to the token embeddings before processing:

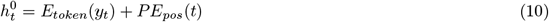

The combination of positional encoding and autoregressive generation enables the model to learn the natural progression of cell-type annotations without requiring explicit hierarchical constraints.

#### Optional Hierarchical Constraint Mechanism

For applications demanding strict adherence to the established cell-type hierarchy ℋ, we implement an optional hierarchical masking mechanism. This mask modifies the output distribution according to:

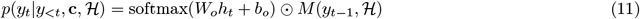

where the mask *M* (*y*_*t*−1_, ℋ) is defined as:

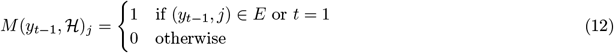

This masking ensures that only valid child terms in the hierarchy can be generated, providing an additional layer of biological consistency when required.

### Training Objective

Lemur’s total loss during pretraining aggregates terms from the encoder and decoder’s learning objectives as seen in:

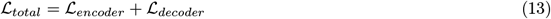

This combined objective enables the model to learn both gene-gene relationships and the natural progression of cell-type annotations from broad to specific categories.

#### Encoder Objective: Masked Language Modeling

The primary encoder pretraining objective is Masked Language Modeling (MLM), initially introduced in natural language processing ^[2]^ and rapidly adapted in single cell foundation models ^[3, 4]^. MLM allows for bidirectional attention by randomly masking input tokens and training the model to predict them based on their surrounding context. However, Lemur performs MLM on both gene tokens and expression values simultaneously and independently, capturing both gene and expression dynamics.

The transcriptomic encoder loss function is as follows:

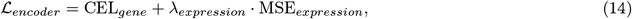

where CEL_*gene*_ represents the Cross Entropy Loss for masked gene token prediction, MSE_*expression*_ is the Mean Squared Error for masked expression value prediction, and *λ*_*expression*_ is a weighting parameter balancing the two objectives. Based on hyperparameter tuning, the masking rates for both gene tokens and expression values were set to 10% (`masking = 0.1`, independent).

#### Decoder Objective: Hierarchical Next Token Prediction

The cell-type decoder’s main pretraining objective is next token prediction. The hierarchical decoder learns in an autoregressive manner, so tokens can only attend to previously generated tokens, enforcing a progression from broader hierarchical terms to specific cell-type labels. For a sequence of cell-type tokens **y** = (*y*_1_, …, *y*_*T*_), the cell-type decoder loss is defined by:

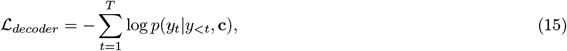

where *y*_*t*_ represents all tokens before position *t*, and **c** is the cell embedding from the transcriptomic encoder serving as the decoder’s memory.

### Training Setup and Optimization

Lemur was trained on three NVIDIA A100 GPUs for approximately two days, completing 10 epochs with a batch size of 64. Training acceleration was achieved using DeepSpeed^[24]^ integrated with Lightning Fabric^[25]^, enabling efficient memory and compute utilization. The training environment leveraged PyTorch version 2.3^[26]^ and FlashAttention-2^[27]^ to optimize attention mechanism speed and memory usage for the large model.

The Adam optimizer with decoupled weight decay regularization (AdamW) was employed with hyperparameters *β*_1_ = 0.9, *β*_2_ = 0.999, *ϵ* = 1 *×* 10^−8^. A cosine learning rate scheduler with a maximum learning rate of 1 *×* 10^−5^ and a linear warmup over the first epoch was used to ensure stable training convergence.

## Data Availability

The model architecture and weights are available on Hugging Face (https://huggingface.co/jorgebotas/lemur). Detailed references to the single-cell and single-nucleus datasets used for training and evaluation are provided in the Methods and Materials section. The lemur icon is freely available on Flaticon.com

---

**Supplementary Figure 1:**
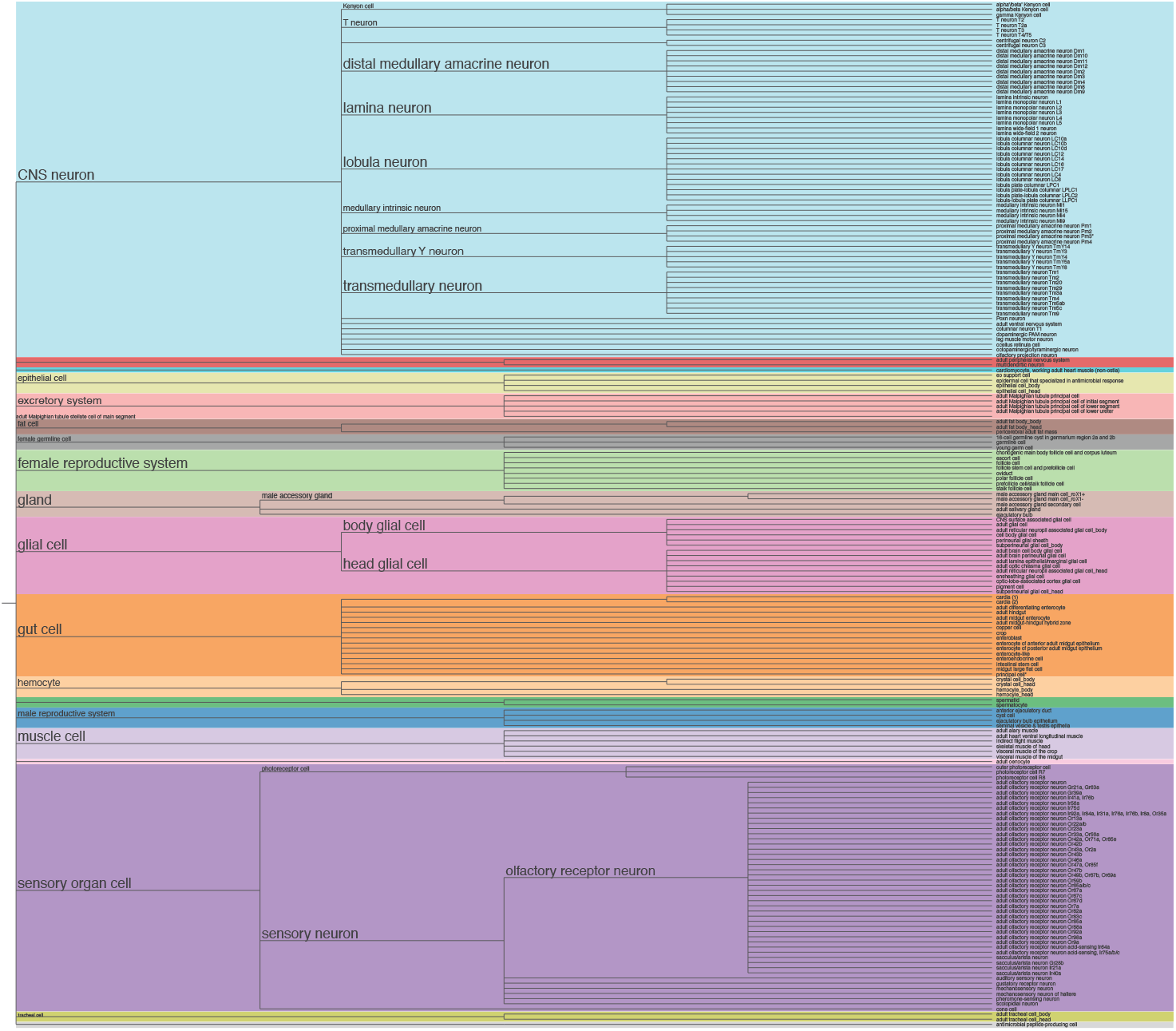
Comprehensive cell type hierarchy used for Lemur’s pretraining. Visualization of the unified cell type ontology derived from the integration of multiple *Drosophila* atlases. The hierarchy comprises 238 distinct annotations organized in a tree structure with 199 leaf nodes and 19 major classes (indicated by different colors). Starting from root nodes (left), the tree progressively branches into increasingly specific cell types (right). Major branches include CNS neurons (light blue), showing extensive subtype specification particularly in the visual system; glial cells (pink) divided into body and head populations; and diverse tissue-specific cell types such as muscle cells (dark blue), sensory neurons (purple), and reproductive system cells (green). This hierarchical organization provides the foundation for Lemur’s generative cell type decoder, enabling consistent predictions across multiple levels of granularity. Tree visualization generated using ETE Toolkit^[12]^.

**Supplementary Figure 2:**
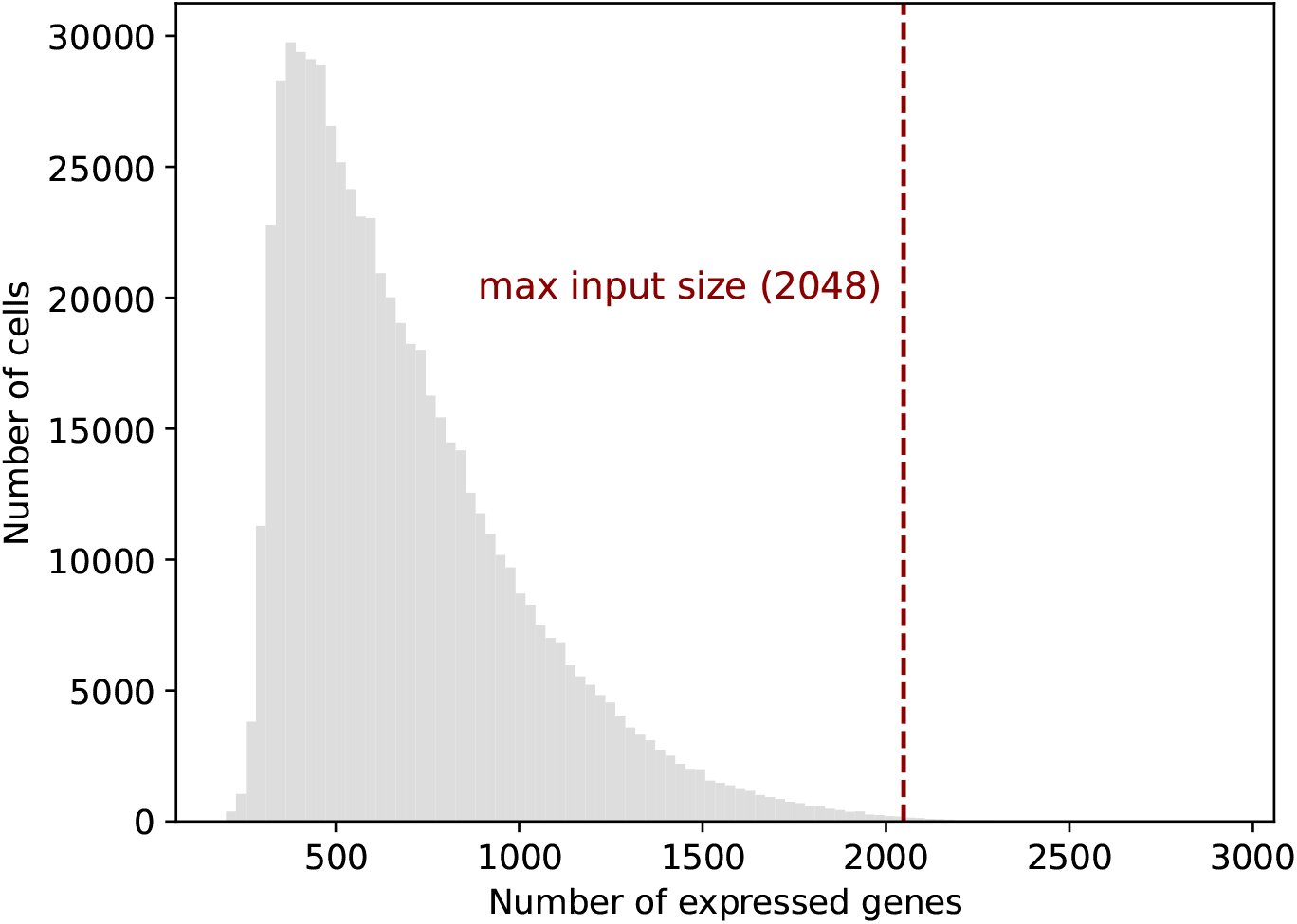
Distribution of gene expression in single-cell transcriptomes used for model pre-training. Histogram showing the number of expressed genes per cell across the complete pre-training dataset, which comprises 624,458 single-cell transcriptomes spanning 16,195 unique genes. The vertical red dashed line indicates Lemur’s maximum input sequence length of 2,048 genes, demonstrating that the model’s architecture can process 99.8% of the cellular transcriptomes in their entirety without truncation. The distribution reveals that most cells express between 400-1,000 genes, with a peak around 500 genes, indicating that Lemur’s input capacity substantially exceeds typical transcriptome complexity. This architectural design ensures comprehensive coverage of gene expression patterns while maintaining computational efficiency.

